# Rare non-coding variants are associated with plasma lipid traits in a founder population

**DOI:** 10.1101/141960

**Authors:** Catherine Igartua, Sahar V Mozaffari, Dan L Nicolae, Carole Ober

## Abstract

Founder populations are ideally suited for studies on the clinical effects of alleles that are rare in general populations but occur at higher frequencies in these isolated populations. Whole genome sequencing in 98 South Dakota Hutterites, a founder population of European descent, and subsequent imputation to the Hutterite pedigree revealed 660,238 single nucleotide polymorphisms (SNPs; 98.9% non-coding) that are rare (<1%) or absent in European populations, but occur at frequencies greater than 1% in the Hutterites. We examined the effects of these rare in European variants on plasma levels of LDL cholesterol (LDL-C), HDL cholesterol (HDL-C), total cholesterol and triglycerides (TG) in 828 Hutterites and applied a Bayesian hierarchical framework to prioritize potentially causal variants based on functional annotations. We identified two novel non-coding rare variants associated with LDL-C (rs17242388 in *LDLR*) and HDL-C (rs189679427 between *GOT2* and *APOOP5*), and replicated previous associations of a splice variant in *APOC3* (rs138326449) with TG and HDL-C. All three variants are at well-replicated loci in genome wide association study (GWAS) but are independent from and have larger effect sizes than the known common variation in these regions. We also identified variants at two novel loci (rs191020975 in *EPHA6* and chr1:224811120 in *CNIH3*) at suggestive levels of significance with LDL-C. Candidate expression quantitative loci (eQTL) analyses in lymphoblastoid cell lines (LCLs) in the Hutterites suggest that these rare non-coding variants are likely to mediate their effects on lipid traits by regulating gene expression. Overall, we provide insights into the mechanisms regulating lipid traits and potentially new therapeutic targets.

## Introduction

Blood lipid traits are under strong genetic control and are modifiable risk factors for cardiovascular disease, one of the leading causes of death^1^. These traits include plasma levels of low-density lipoprotein cholesterol (LDL-C), high-density lipoprotein cholesterol (HDL-C), total cholesterol and triglycerides (TG) and have estimated heritabilities of 40-60% across populations^2–4^. Although genome-wide association studies (GWAS) of lipid traits have been successful in identifying hundreds of common variants with robust associations, the associated variants account for only about 10-14% of the total phenotypic variance^5,6^. While a portion of the unexplained genetic variance may result from overestimates of heritability and complex genetic architectures, such as those involving epistasis or gene-environment interactions^7^, the effect of rare loss-of-function variation in complex traits is understudied and likely contributes to the heritability of blood lipid traits and risk for cardiovascular disease.

In fact, sequencing studies in families or patients with rare monogenic lipid disorders have uncovered many novel genes harboring rare coding mutations of large effect and revealed critical pathways for lipid metabolism^8–10^. These studies have supported earlier observations suggesting that rare variants in the general population contribute significantly to lipid traits and possibly more generally to common, complex phenotypes. For example, a resequencing study of genes that harbor causal mutations in monogenic lipid disorders identified an enrichment of nonsynonymous variants associated with lipid levels by sequencing unrelated individuals sampled from the tails of the HDL-C trait distribution^11^. This study demonstrated for the first time that rare variants contribute to population-level variation in blood lipids.

While providing valuable insight into the mechanisms regulating lipid traits, rare variant studies in complex traits have focused on coding regions of the genome, largely due to the recent explosion of exome sequencing studies and the relative ease in interpreting these findings^12^. As a result, the non-coding portion of the genome has been largely unexplored. Similar to common non-coding variants, rare non-coding variants may also impact gene expression and protein abundance, but the sample sizes of most studies of gene expression are underpowered to identify the independent effects of rare variants. A few studies have provided strong support for the aggregated effect of multiple rare non-coding variants on nearby gene expression, and identified enrichments of rare non-coding variation for individuals showing extreme gene expression levels, as compared with the same genes in non-outlier individuals^13–15^, and shown cis eQTL effect sizes are significantly higher for SNPs with lower allele freuquencies^16^. Despite this evidence, the broader questions of what functional features characterize rare non-coding variants influencing gene expression and how different functional classes of rare non-coding variations influence disease are unknown.

Founder populations offer the opportunity to study the effects of variants that are rare in general populations but have reached higher frequencies in these isolated populations due to the effects of random genetic drift in the early generations after their founding^17–19^. In addition, their overall reduced genetic complexity and relatively homogenous environments and lifestyles can enhance the effects of rare genetic variants on phenotypic traits and thereby facilitate the detection of susceptibility loci that underlie complex disease, as elegantly illustrated in studies of the Amish and Icelandic populations^20–22^.

In this study, we dissect the genetic architecture of plasma lipid traits in members of the South Dakota Hutterites, a founder population of European descent. The 828 individuals who participated in these studies are related to each other in a 13-generation pedigree with 64 founders. We performed GWAS using 660,238 “rare in European variants” (REVs) that occur at frequencies greater than 1% in the Hutterites, and integrated functional and regulatory annotations that allowed us to narrow down potential candidate variants despite the long-range linkage disequilibrium present in the population. Our studies revealed rare variants with large effects at two novel and three known blood lipid loci associated with LDL-C, HDL-C or TG, potentially yielding novel insights into the mechanisms regulating lipid traits and new therapeutic targets.

## Results

Approximately 7 million single nucleotide polymorphisms (SNPs) identified through whole genome sequencing in 98 Hutterite individuals were imputed using the known identity by descent (IBD) structure of the Hutterite pedigree^23^ to the 828 individual in our study. We selected 660,238 variants that were either absent or rare (<1%) in European databases (see Methods) and occurred at frequencies greater than 1% in the Hutterites (REVs) for association testing with fasting lipid measurements (plasma LDL-C, HDL-C, total cholesterol and TG levels). Based on their RefSeq^24^ annotations, the majority of these REVs were intergenic (54.2%) or intronic (43.7%); the remaining 2.1% were exonic (1%), in the 3’ or 5’ UTR (1.1%) or predicted to affect splicing (6.4×10^−5^%; **Figure 1**).

**Figure 1:**
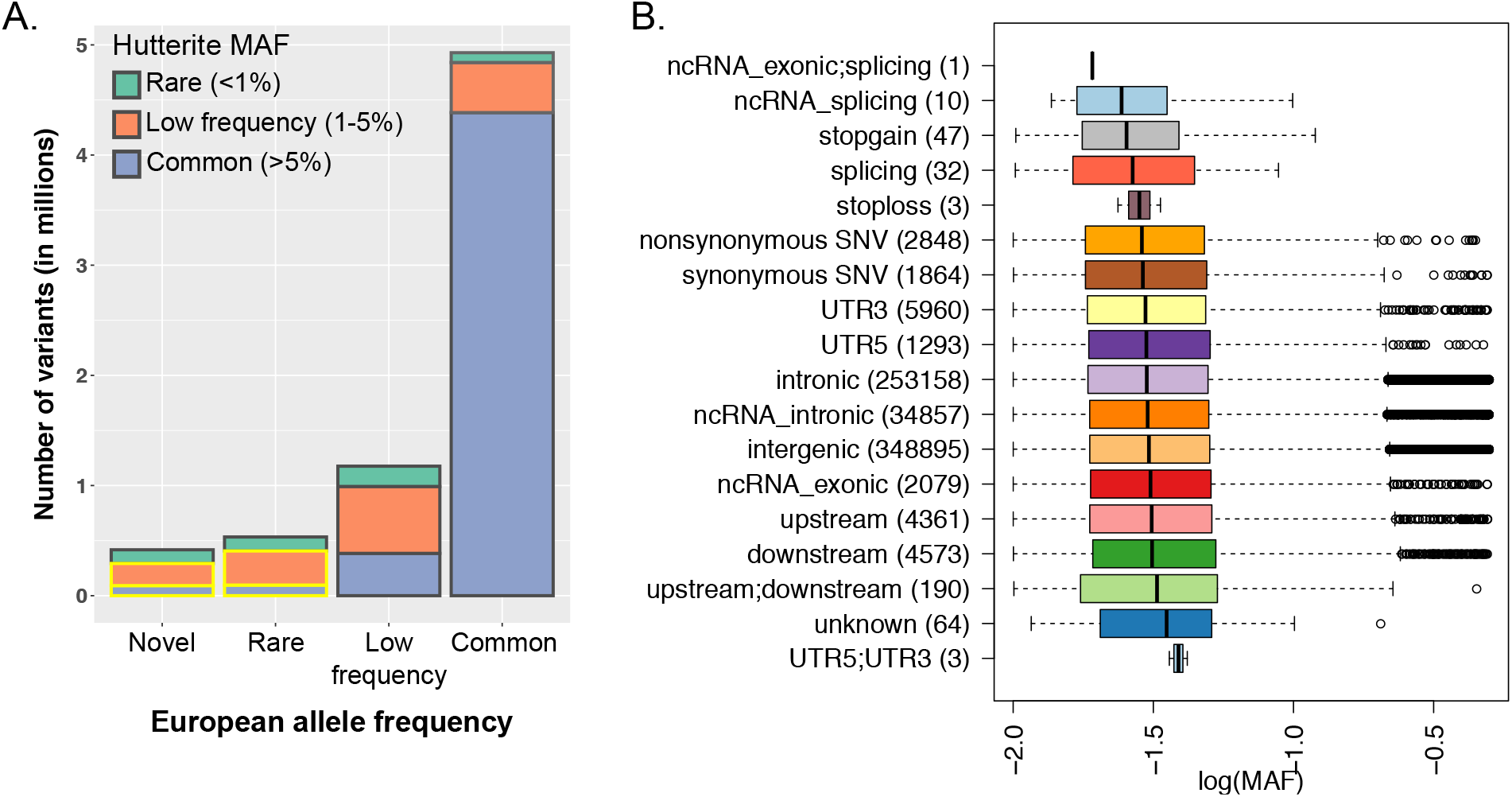
Allele frequencies of variants present in Hutterite genomes. (A) *Bar plot of variants binned by maximum reported allele frequency in European databases, stratified by European minor allele frequency (MAF) on the x-axis and shown by Hutterite minor allele frequency within each bar*. Variants presented include ~7 million variants discovered in 98 Hutterite whole genome sequences^23^. Maximum European allele frequencies were calculated from surveys of the Exome Sequencing Project (ESP), Exome Aggregation Consortium (ExAC) and the 1000 genomes project. Rare in European variants (REVs) included in the lipid trait association studies are highlighted with a yellow border in the novel and rare categories. (B) *Boxplots of log(MAF) by annotation category for the 660,238 REVs included in the lipid trait association studies*. Annotation categories were based on RefSeq^24^. Numbers in parenthesis correspond to the number of variants in each annotation class. Black vertical lines correspond to the median and whiskers to the 25^th^ and 75^th^ percentiles.

### Single Variant GWAS

For each lipid trait, we performed association analyses in two stages. First, we performed single variant analyses using a linear mixed model (GEMMA^25^), including age and sex as fixed effects and kinship as a random effect. Quantile-quantile plots showed no inflation of test statistics for any of the lipid traits (**Figure S1**); the most significant associations at each loci are presented in **Table 2** (Manhattan plots for all GWAS are shown in **Figure S2**). We detected two genome-wide significant loci (p<5×10^−8^) associated with either increased LDL-C or with reduced TG levels and contained multiple rare variants of large effect in regions previously associated with lipid traits in GWAS^5,6^. These two loci include 78 REVs associated with increased LDL-C over a 7.6 Mb region flanking the LDL receptor gene (*LDLR*) on chromosome 19, and 39 REVs associated with reduced TG levels over a 3.8 Mb region flanking the Apolipoprotein C3 (*APOC3*) gene on chromosome 11. The latter includes a known rare splicing variant in *APOC3* (rs138326449)^26,27^. No genome-wide significant associations with REVs were detected for HDL-C or total cholesterol.

**Table 1:** Sample composition and clinical descriptions.

| Lipid trait | Sample size<br>(Male/Female) | Mean age<br>[95% range] | Mean mg/dL<br>[95% range] |
| --- | --- | --- | --- |
| Low density lipoprotein cholesterol (LDL-C) | 807 (367/440) | 36.04<br>[16-65] | 125.1<br>[48.86-201.34] |
| High density lipoprotein cholesterol (HDL-C) | 828 (381/447) | 36.35<br>[16-66] | 54.7<br>[23.78- 85.62] |
| Triglycerides (TG) | 828 (382/446) | 36.31<br>[16-65.65] | 121.5<br>[0-299.91] |
| Total cholesterol | 828 (381/447) | 36.35<br>[16-66] | 203.9<br>[121.8-286] |

**Table 2:** Genome-wide significant results for plasma lipid trait GWAS of rare in European variants. Variants presented are the most significant in each region.

| Trait | rsID<br>(chr:location) | Minor/<br>major | MAF | 1000G<br>EUR AF | p | Beta<br>(SE) | Variant type<br>(gene) | No. of<br>significant REVs |
| --- | --- | --- | --- | --- | --- | --- | --- | --- |
| LDL-C | rs557778817<br>19:11305534 | T/C | 0.032 | 0.003 | $1.48 \times 10^{-17}$ | 1.30<br>(0.15) | intronic<br>(*KANK2*) | 78 |
| TG | rs184333869<br>11:117947268 | T/C | 0.023 | 0.001 | $5.41 \times 10^{-13}$ | -1.28<br>(0.17) | ncRNA intronic<br>(*TMPRSS4-AS1*) | 39 |

rsID was annotated with dbSNP142. Location is based on hg19. Direction of effect corresponds to the minor allele.

### Application of fgwas to lipid traits

In second stage analyses, we annotated all variants discovered in the Hutterites using 25 sequence-based annotations and 332 functional annotations (see Methods) and applied the functional GWAS (fgwas)^28^ framework to further evaluate the GWAS results and prioritize candidate rare variants based on prior functional knowledge. We split the genome into blocks averaging 125Kb (~50 REVs) and performed forward selection to build models that combined effects from multiple annotations followed by a cross validation step to avoid overfitting while maximizing the likelihood of each model. **Figure 2** shows the maximum likelihood estimates (MLE) and 95% CI of the enrichment effects for the selected annotations in each of the final joint models for each of the blood lipid traits. Annotation descriptions and the respective penalized effects used in each model are provided in **Supplemental Tables 1-4**.

**Figure 2:**
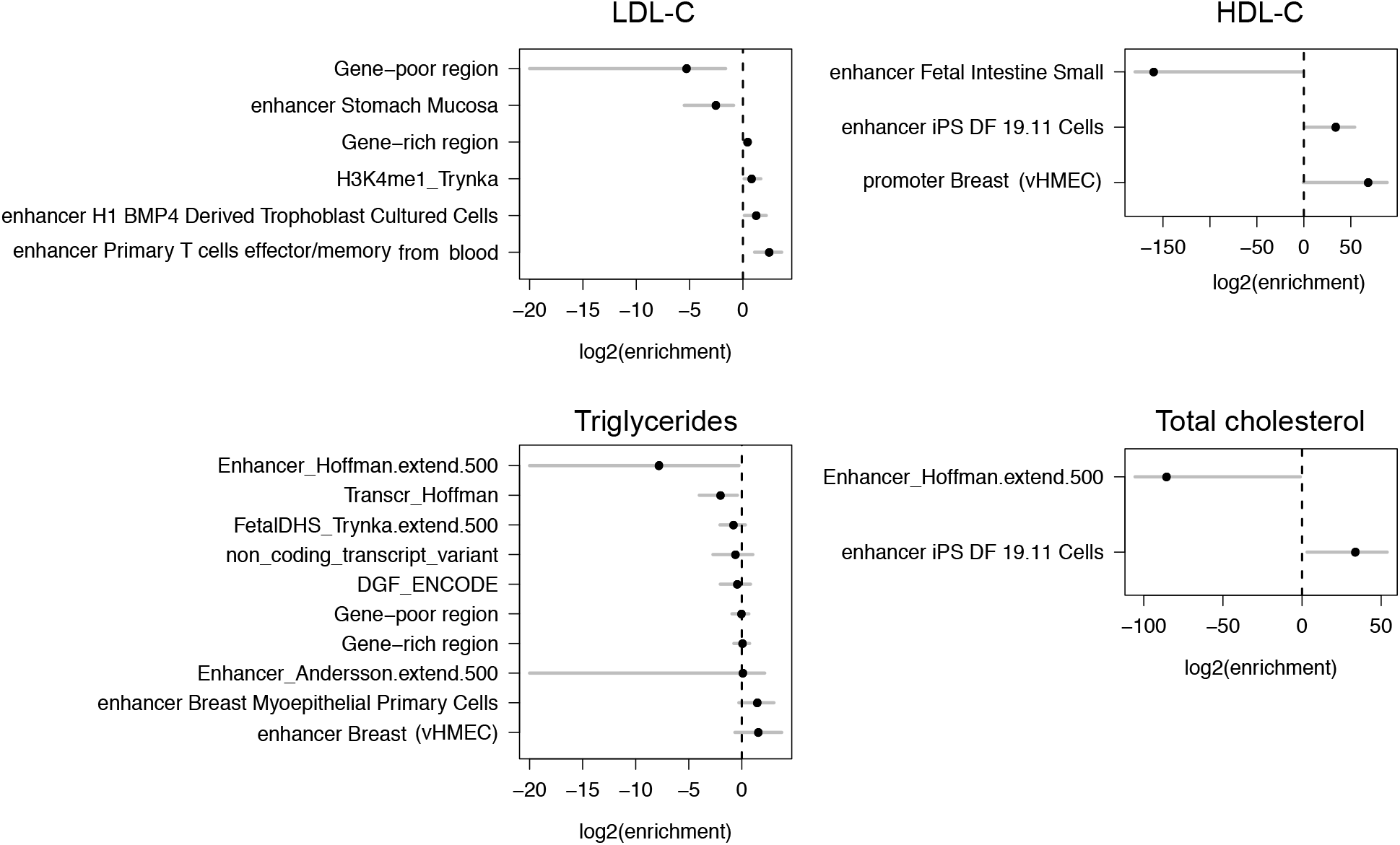
Joint fgwas models for blood lipid traits. Models were fit combining multiple annotations using the fgwas hierarchical methodology, as described in Pickrell *et al*. (2014)^28^. The MLE and 95% CI of the enrichment effects of each annotation in the final model (modeling performed using penalized likelihood) are shown.

The joint model for each fgwas estimated both the probability that each block contains a causal variant and the posterior probability that a variant is causal conditional on the presence of one causal variant in the region. Variants with the largest posterior probabilities of causality will tend to have the most significant p-values and functional annotations that predict association genome-wide. We weighted our GWAS results based on the fgwas joint model and applied a regional prior probability of association (PPA) threshold of 0.9. Overall, we identified 63 consecutive blocks associated with LDL-C on chromosome 19 and 20 blocks on chromosome 11 associated with TG, all of which harbor REVs that passed genome-wide significance in the single variant GWAS (p<5×10^−8^), and additional blocks associated with HDL-C in a novel region on chromosome 16 and one on chromosome 11. In addition, when we applied a slightly lower regional threshold of PPA>0.75, additional blocks associated with LDL-C on chromosomes 1 and 3 were identified. We present all variants within regions with PPA>0.75 and SNP PPA>0.5 in **Table S5** and refer to these as candidate variants.

#### LDL-C

The GWAS of LDL-C identified 78 genome-significant REVs associated with increased levels of LDL-C on a haplotype that spans 7.6Mb on chromosome 19p13.2. Despite the high LD and resulting long haplotype, we were able to prioritize 35 candidate variants with SNP posterior probabilities greater than 0.75, 13 of which had posterior probabilities equal to one (PPA=1; Table S5). Among these 13 variants, one satisfied all the annotations selected in the model. This is a SNP located in the first intron on the *LDLR* (rs17242388; logBF=28.02; p_gwas_=3.85×10^−15^; MAF=3%; 1000 genomes EUR MAF=0.6%). While the established function of the *LDLR* in lipid metabolism makes an intronic LDLR variant an obvious candidate^32^, a novel nonsynonymous variant in Zinc Finger Protein 439 (*ZNF439;* chr19:11978399-T) was the fgwas candidate variant with the highest predictive functional score (CADD^33^=16.65; PPA=0.8; logBF=20.95; p_gwas_=5.55×10^−12^; MAF=0.04).

We identified two additional loci that were suggestively associated with LDL-C (regional PPA>0.75). The first association was a protective variant private to the Hutterites located in the first intron of the Cornichon Family AMPA Receptor Auxiliary Protein 3 gene (*CNIH3*) and was associated with decreased LDL-C (chr1:224811120; logBF=5.26; PPA=0.76; p_gwas_=7.99×10^−5^; MAF=0.02). The second variant was an intronic variant on chromosome 3 in the EPH Receptor A6 gene (*EPHA6;* rs191020975) associated with increased LDL-C (logBF=6.46; PPA=0.52; pgwas=1.88×10^−5^; MAF=0.02; 1000 genomes EUR MAF=0.001).

#### Triglycerides

Out of the 39 REVs on a 3.8Mb haplotype associated with TG levels, 14 were potentially causal with SNP posterior probabilities of association greater than 0.75. The SNP with the highest posterior probability (rs149157643; PPA=1; logBF=22.49; p_gwas_=7.47×10^−13^; MAF=2.38%; 1000 genomes EUR MAF=0.7%) is an intronic variant within the non-coding RNA (ncRNA) gene Transmembrane Protease Serine 4 Antisense RNA 1 (*TMPRSS4-AS1*) and is 25Kb away from the most associated variant in the region (rs184333869; PPA=0.97; p=5.41×10^−13^; MAF=2.40%; 1000 genomes EUR MAF=0.1%). Chromosome 11q23 is a well replicated GWAS locus for multiple lipid traits^5,34^ (**Figure 4**) with several implicated rare variants, including loss-of-function mutations in *APOC3*^20,26,27^ associated with decreased TG levels. In fact, the candidate variant associated with TG in the Hutterites with the highest functional and conservation score in this region (CADD=25.1; GERP=4.89) is a previously reported splice variant in *APOC3* (rs138326449; MAF=2.23%; 1000 genomes MAF=0.3%) but had a slightly lower posterior probability of association in our model (PPA=0.86; logBF=15.38; p_gwas_=1.08×10^−9^) compared to other variants in the region. To our knowledge, the role of this potential splice donor *APOC3* polymorphism (rs138326449) in regulation of plasma lipids has not been characterized thus far. Therefore, while there is be compelling evidence that reduced plasma levels of APOC3 protein results in lower TG levels^20,26^, there may be multiple rare variants within an extended haplotype influencing TG levels in the Hutterites.

**Figure 3:**
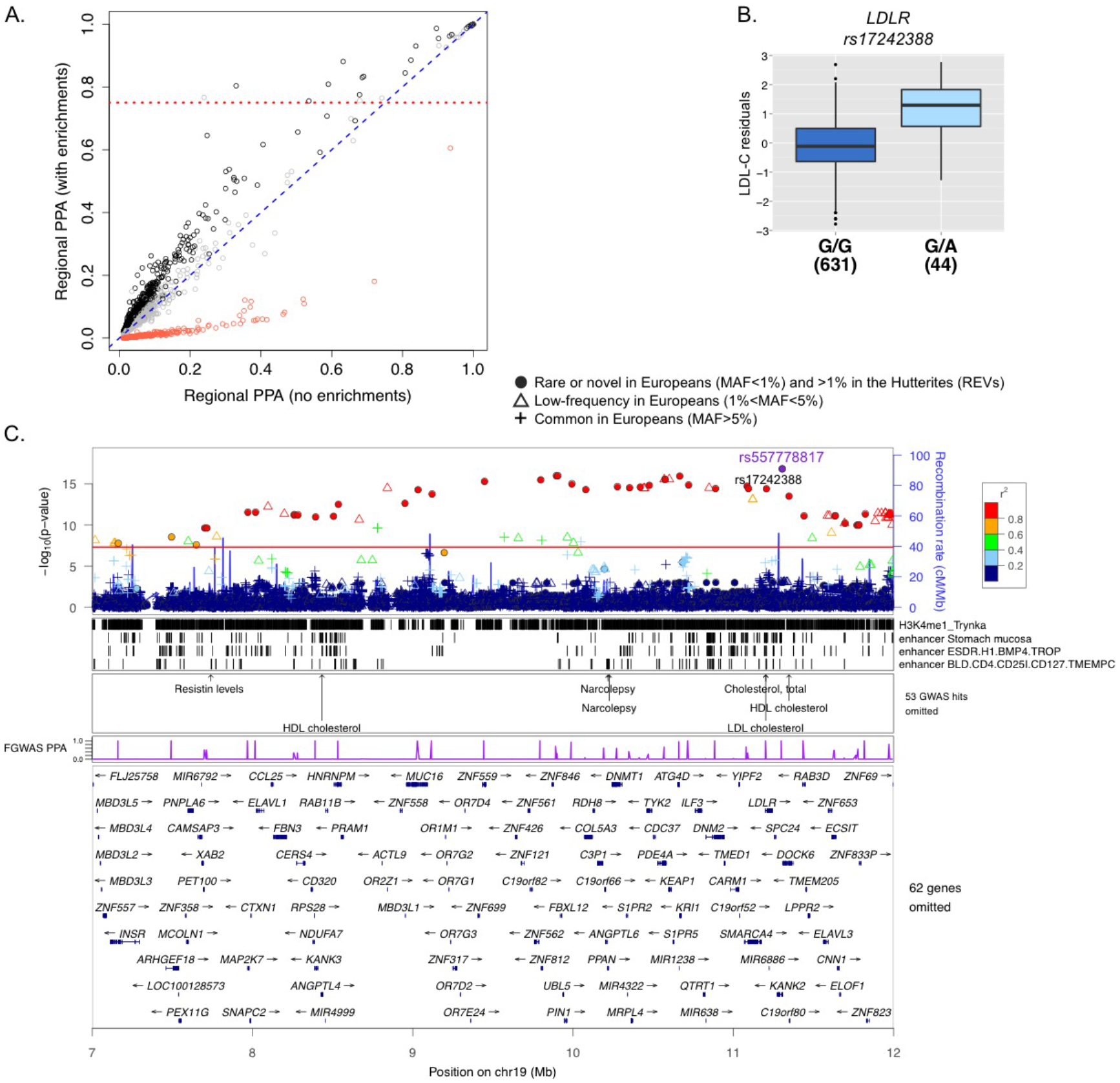
Rare variants on chromosome 19 are associated with LDL-C. (A) *Reweighted GWAS*. GWAS results were reweighted using the joint model presented in **Figure 2**. Each point represents a region of the genome and its corresponding posterior probability of association (PPA) in a model with and without enrichments. Red points correspond to regions in the bottom, grey points in the middle, and black points in the top tertiles of gene density. The dotted red line represents regional PPA > 0.75 in the enriched model. (B) *Boxplot of association between LDL-C (y-axis) and candidate LDLR expression level by variant rs17242388 (p=3.89×10^−15^) (x-axis)*. Black horizontal lines show medians and whiskers show the 25^th^ and 75^th^ percentiles. (C) *Locus plot*. The top panel shows the p-values of association with LDL-C for all variants discovered in the Hutterites regardless of allele frequency in Europeans (y-axis). Symbols correspond to the maximum allele frequency in Europeans, with closed circles representing REVs (see legend), and are colored based on their LD r^2^ with the most associated variant in the region (rs557778817). The next three panels show tracks for the annotations selected in the fgwas joint model, annotations of known GWAS loci from NHGRI, and the estimated PPA of being causal for each variant in the reweighted fgwas. The bottom-most panel shows the genes in the region.

**Figure 4:**
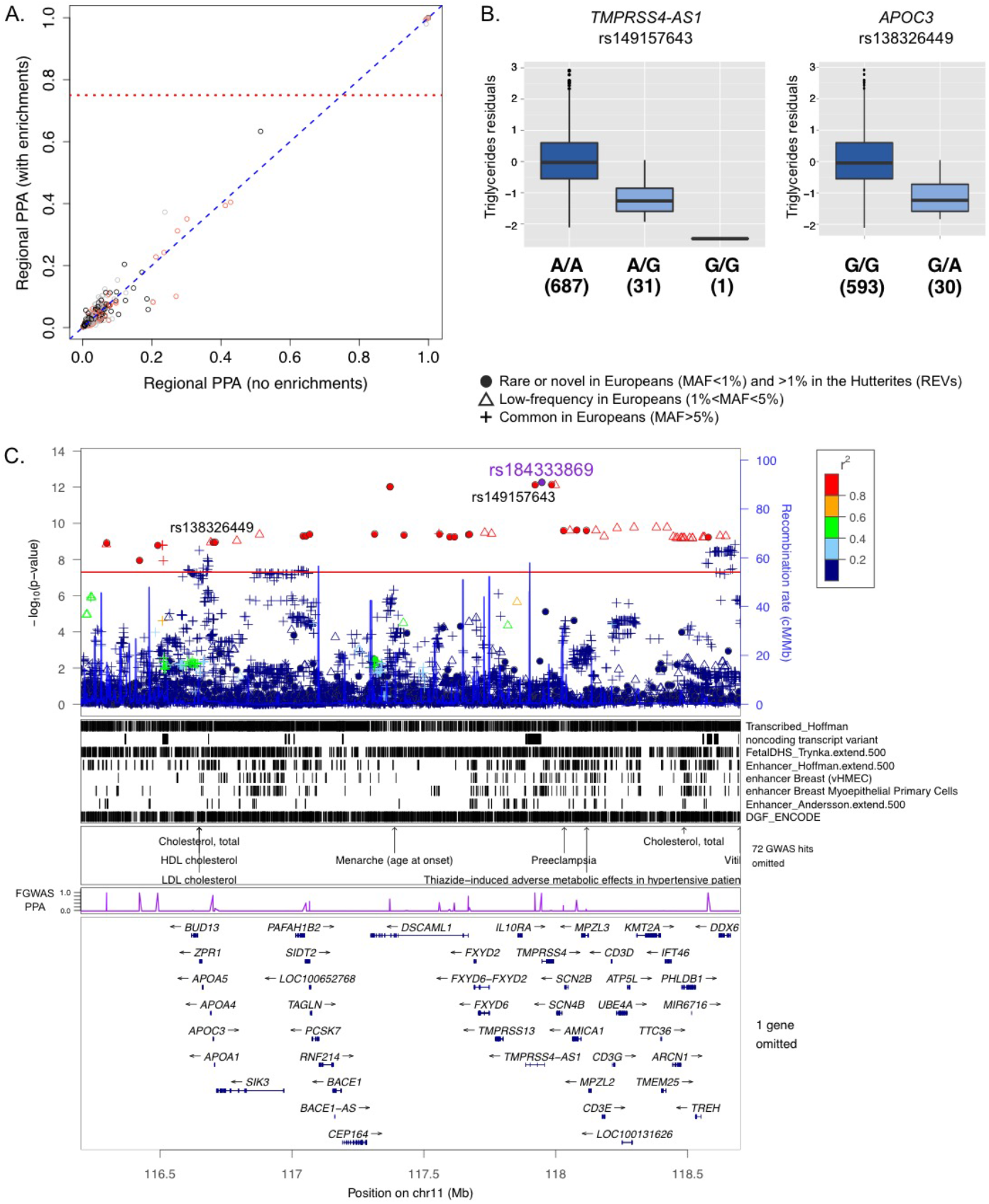
Rare variants on chromosome 11 are associated with TG levels. (**A**) *Reweighted GWAS*. GWAS results were reweighted using the joint model presented in **Figure 2**. Each point represents a region of the genome and its corresponding posterior probability of association (PPA) in a model with and without enrichments. Red points correspond to regions in the bottom tertile of gene density, grey points the middle and black points the top. Dotted red line represents regional PPA > 0.75 in the enriched model. (**B**) *Boxplots of association between TG (y-axis) and candidate noncoding RNA intronic variant in TMPRSS4-AS1 (rs149157643; p=7.47×10^−13^) and splicing variant in APOC3 (rs138326449; p=1.08×10^−9^)*. Black horizontal lines show medians and whiskers show the 25^th^ and 75^th^ percentiles. (**C**) *Locus plot*. The top panel shows the p-values of association with TG for all variants discovered in the Hutterites regardless of allele frequency in Europeans. Symbols correspond to the maximum allele frequency in Europeans, with closed circles representing REVs (see legend), and are colored based on their LD r^2^ with the most associated variant in the region (rs184333869). The next three panels show tracks for the annotations selected in the fgwas joint model, annotations of known GWAS loci from NHGRI, and the estimated PPA of being causal for each variant in the reweighted fgwas. The bottom-most panel shows the genes in the region.

#### HDL

Although the HDL-C GWAS did not identify any genome-wide significant REVs, the fgwas revealed two loci associated with HDL-C with regional PPA greater than 90%. The first locus was associated with increased HDL-C levels and tags the same haplotype on chromosome 11q23.3 that is associated with lower TG levels. The selected variant at this locus with the highest probability was also the most significant SNP in the HDL-C GWAS (rs184333869; logBF=9.03; PPA=0.86; p_gwas_=1.1×10^−6^; **Figure 5**) and the most significant association in the TG GWAS. The second association was with variants in a 3.8Mb region on 16q13 and decreased HDL-C levels; this is one of the most replicated loci for HDL-C and cardiovascular disease risk^5,35^ (rs3764261, a variant upstream of Cholesteryl Ester Transfer Protein [CETP]). The variant with the highest posterior probability of association was rs189679427 (PPA=0.76; logBF=6.58; p_gwas_=1.27×10^−6^; MAF=5.2%; 1000 genomes EUR MAF=0.2%; **Table S5**), an intergenic variant located 143Kb from Glutamic-Oxaloacetic Transaminase 2 (*GOT2*) and 877Kb from Apolipoprotein O Pseudogene 5 (*APOOP5*). While *GOT2* has not been directly linked to HDL-C levels, its characterized function as a membrane associated fatty acid transporter highly expressed in the liver, the primary tissue for apolipoprotein metabolism^22,36,37^, makes it an interesting candidate.

**Figure 5:**
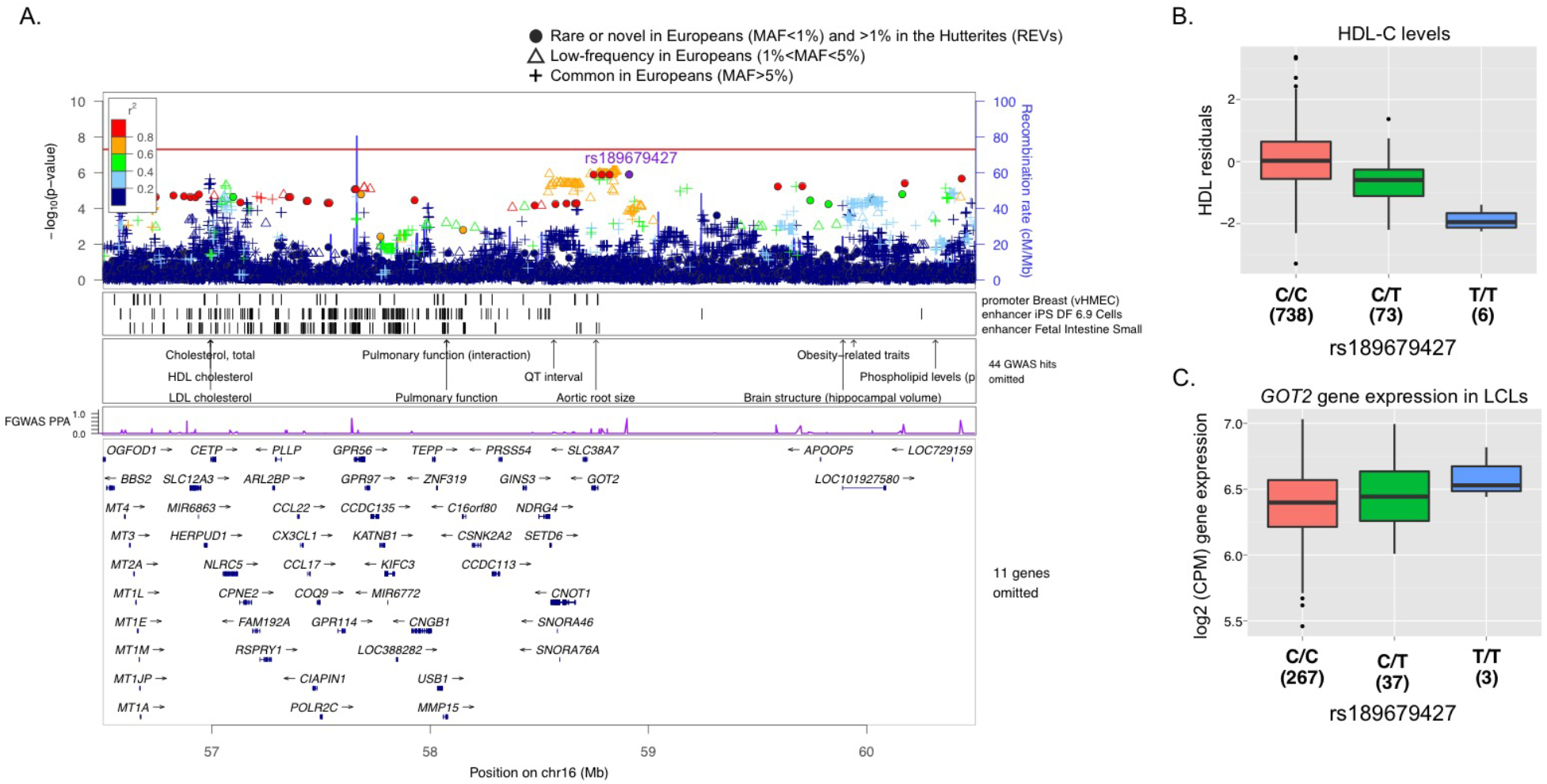
Rare variants on chromosomes 16 are associated with HDL-C. (***A***) *Locus plots*. The top panel shows the p-values of association with HDL-C for all variants discovered in the Hutterites regardless of allele frequency in Europeans. Symbols correspond to the maximum allele frequency in Europeans, with closed circles representing REVs (see legend), and are colored based on their LD r^2^ with the most associated variant in the region (rs189679427). The next three panels show tracks for the annotations selected in the fgwas joint model, annotations of known GWAS loci from NHGRI, and the estimated PPA of being causal for each variant in the reweighted fgwas. The bottom-most panel shows the genes in the region. (**B**) Boxplots of association between *HDL-C levels (y-axis) and* rs189679427 (*intergenic variant between GOT2 and APOOP5; p=1.27×10^−6^*). (**C**) Genotype boxplots of rs189679427 *eQTL boxplots for GOT2 expression in LCLs* (*p=0.004*). Numbers beneath genotypes correspond to the number of individuals in each class.

### Conditional analyses and candidate expression quantitative trait locus (eQTL) studies in the Hutterites

We performed two sets of conditional analyses for each trait with one or more significant associations (LDL-C, TG and HDL-C), including all variants present in the Hutterites genomes that resided each of the associated regions regardless of their minor allele frequencies in Europeans. First, we conditioned on the most significant rare variant in our analyses to assess whether other (rare or common) variants in the region either contribute to the observed association signal or are independently associated with the trait but whose effects were masked by the larger effect of the rare variant. Second, in regions with known associated variants from other GWAS, we also conditioned on the GWAS variant(s) to verify that the rare variant signal in our study is independent of known associations at this locus (**Table 3**). We then evaluated the evidence for regulatory effects of the candidate variants (**Table S5**) on genes within 250Kb of the variants using gene expression data in lymphoblastoid cell lines (LCLs) from the Hutterites^38^. Although LCLs have known limitations, genetic effects on gene expression are often shared across multiple tissues^15,39^. Importantly, however, our focus here on private or rare variation in the Hutterites makes it impossible to utilize publicly available eQTL databases in other tissues to assess our candidate variants.

**Table 3:** Summary results for conditional analyses. (**A**) *Conditional analysis on lead rare variant identified in this study*. For each candidate loci, the listed variant represents the next lead variant in the region (MAF>1%) after regressing the effect of the lead rare variant identified each GWAS. (**B**) *Conditional analysis on previously reported common GWAS variants in the region*. Candidate causal REVs in GWAS loci remain significant after taking into account the effects from known common variation.

| **(A) Conditional analysis on lead rare variants** |  |  |  |  |  |  |  |  |
| --- | --- | --- | --- | --- | --- | --- | --- | --- |
| Trait | Conditioning REV ($p_{\text{gwas}}$) | rsID (chr:location) | Minor/major | MAF | $p_{\text{gwas}}$ | $p_{\text{conditional}}$ | Beta<sub>conditional</sub> (SE) | Variant type (gene) |
| LDL-C | rs557778817<br>($p=1.48 \times 10^{-17}$) | rs513663<br>(19: 10539537) | G/A | 0.282 | 0.001 | $3.03 \times 10^{-6}$ | 0.26<br>(0.06) | intronic<br>(PDE4A) |
| LCL-C | chr1:<br>224811120<br>($p=7.99 \times 10^{-5}$) | chr1:224894756 | T/C | 0.03 | 0.0002 | 0.0003 | 0.57<br>(0.16) | intronic<br>(CNIH3) |
| LDL-C | rs191020975<br>($p=1.88 \times 10^{-5}$) | rs538668869<br>(3:96038119) | T/C | 0.01 | 0.02 | 0.01 | 0.50<br>(0.20) | intergenic<br>(MTHFD2P1, MIR8060) |
| TG | rs184333869<br>($p=5.41 \times 10^{-13}$) | rs11604424<sup>a</sup><br>(11: 116651115) | C/T | 0.225 | 0.0002 | $9.50 \times 10^{-9}$ | 0.36<br>(0.06) | downstream<br>(ZPR1) |
| HDL-C | rs184333869<br>($p=1.1 \times 10^{-6}$) | rs56107015<br>(11:121083121) | G/A | 0.170 | $3.74 \times 10^{-5}$ | 0.001 | 0.28<br>(0.06) | Intergenic<br>(TECTA, SC5D) |
| HDL-C | rs189679427<br>($p=1.27 \times 10^{-6}$) | rs4783961<sup>a</sup><br>(16:56994894) | G/A | 0.468 | 0.002 | $1.66 \times 10^{-5}$ | -0.26<br>(0.06) | upstream<br>(CETP) |
| **(B) Conditional analysis on previously reported GWAS variants** |  |  |  |  |  |  |  |  |
| Trait | Conditioning variant ($p_{\text{gwas}}$) | Candidate variant | Minor/major | MAF | $p_{\text{gwas}}$ | $p_{\text{conditional}}$ | Beta<sub>conditional</sub> (SE) | Variant type (gene) |
| LDL-C | rs6511720<sup>a</sup><br>($p=0.009$) | rs17242388<br>(19:11206861) | A/G | 0.03 | $3.89 \times 10^{-15}$ | $1.31 \times 10^{-15}$ | 1.27<br>(0.16) | intronic<br>(LDLR) |
| TG | rs964184<sup>a,b</sup><br>($p=3.13 \times 10^{-9}$) | rs138326449<br>(11:116701354) | A/G | 0.022 | $1.08 \times 10^{-9}$ | $9.85 \times 10^{-9}$ | -1.05<br>(0.18) | splicing<br>(APOC3) |
| HDL-C | rs964184<sup>a</sup><br>($p=0.2$) | rs184333869<br>(11:117947268) | T/C | 0.024 | $1.10 \times 10^{-6}$ | $1.63 \times 10^{-6}$ | 0.88<br>(0.18) | ncRNA intronic<br>(TMPRSS4-AS1) |
| HDL-C | rs17231506<sup>a</sup><br>($p=3.85 \times 10^{-6}$) | rs189679427<br>(16:58911464) | T/C | 0.052 | $1.27 \times 10^{-6}$ | $5.93 \times 10^{-5}$ | -0.54<br>(0.13) | intergenic<br>(GOT2, APOOP5) |
| rsID was annotated with dbSNP142. Direction of effect corresponds to minor allele. Superscripts on rsIDs correspond to study where variant was reported as significant. <sup>a</sup>Global Lipids Genetics Consortium *et al.* Discovery and refinement of loci associated with lipid levels. *Nat. Genet.* **45**, 1274–1283 (2013). <sup>b</sup>Teslovich, T. M. *et al.* Biological, clinical and population relevance of 95 loci for blood lipids. *Nature* **466**, 707–713 (2010). |  |  |  |  |  |  |  |  |

Conditioning on the lead rare variant on chromosome 19 that is associated with LDL-C (rs557778817) revealed a novel common variant in the third intron of Phosphodiesterase 4A (*PDE4A;* rs513663) that reached suggestive significance after removing the effects of the rare variant (p_gwas_=0.001; p_conditional_=3.0×10^−6^; **Table 3** and **Figure 6A**). *PDE4A* plays a key role in many physiological process by regulating levels of the cAMP, a mediator of response to extracellular signals^40^, but to our knowledge variation with this variant or any variants in LD with it has not been previously linked to lipid traits. Both the *PDE4A* common (rs513663) and the *LDLR* rare variant (rs1724388) are associated with higher LDL-C but reside on different haplotypes in the Hutterites, with independent and additive effects of the minor alleles both on lowering *LDLR* gene expression (p=0.008) and increasing plasma levels of LDL-C (**Figure 6B**; p=2.4×10^−8^). We also performed conditional analysis with a commonly replicated variant in the *LDLR* gene that is associated with decreased LDL-C levels and lower risk for coronary heart disease (CHD)^5,6,41^ (rs6511720; p_gwas_=0.009 in the Hutterites) and confirmed that the identified rare variants in the Hutterites have independent and opposite effects compared to the common GWAS variant at this locus (**Table 3**).

**Figure 6:**
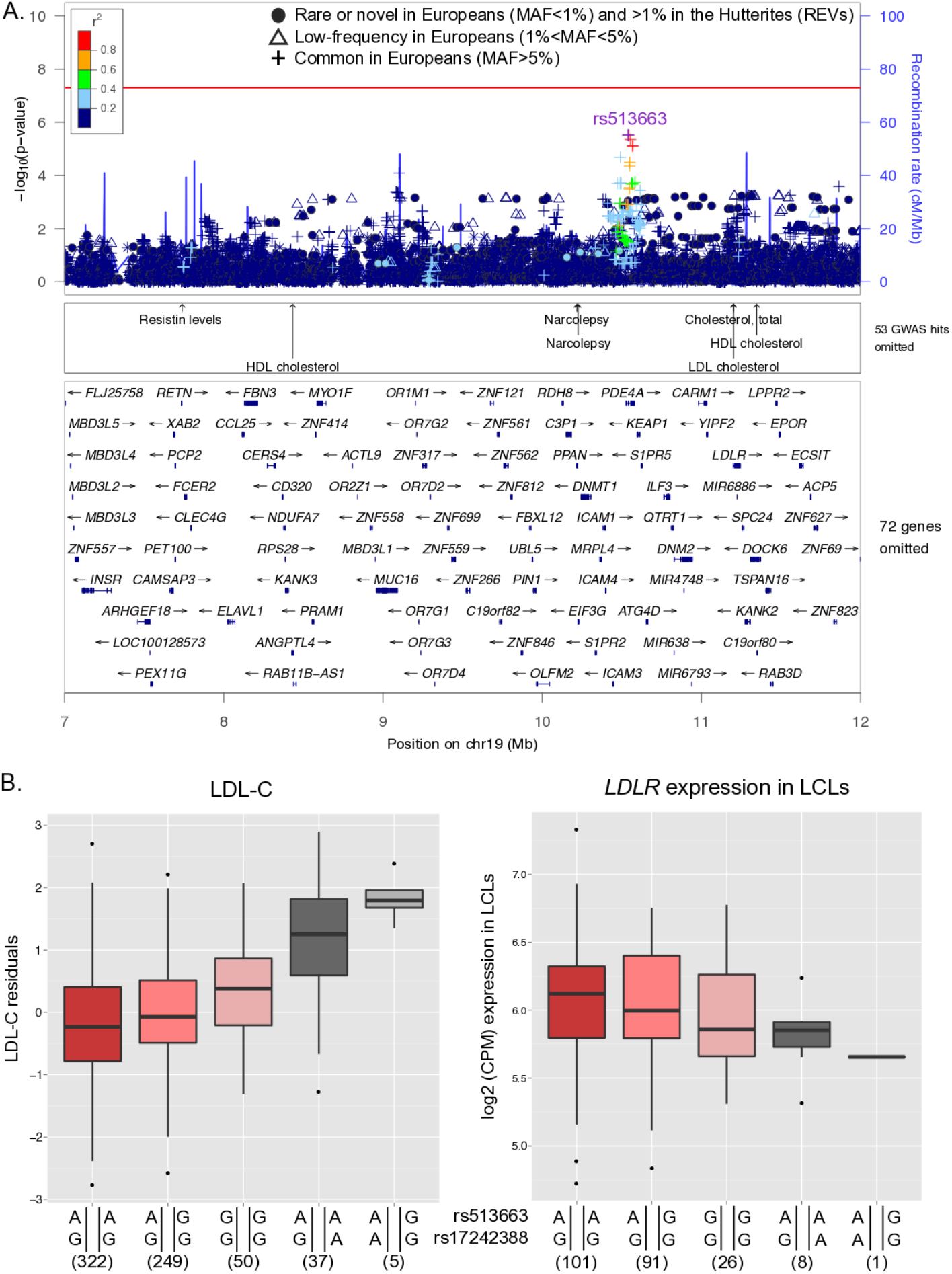
LDL-C conditional analysis on top rare variant (rs557778817). (**A**) *Locus plot*. Conditional analysis on rs557778817 identified novel common variants with suggestive significance. The next three panels show annotations of known GWAS loci from NHGRI and genes in the region. (**B**) *LDLR and PDE4A variant haplotype boxplots for LDL-C levels and LDLR LCL expression*. Phased alleles are the lead common signal in *PDE4D* identified in conditional analysis (rs513663; top) and the candidate *LDLR* rare intronic variant identified in the fgwas (rs17242388; bottom). The trends observed between each of five haplotype combinations present in our sample suggest these variants have additive effects that lower *LDLR* expression and increase LDL-C levels.

We performed eQTL analyses with the 53 LDL-C candidate variants on chromosome 19 (SNP PPA>0.5) and found three out of 245 genes tested, Zinc Finger Protein 440 (*ZNF440*), Dihydrouridine Synthase 3 Like (*DUS3L*) and Hook Microtubule Tethering Protein 2 (*HOOK2*), had expression levels associated with at least one candidate variant at a p<10^−4^ (**Table S5**). The most significant eQTL in this region was a private nonsynonymous variant on *ZNF439* (chr19:11978399) and decreased levels of Zink Finger Protein 440 (*ZNF440;* p=2×10^−11^).

Conditional analyses of the lead rare variant on chromosome 11 (rs184333869) that was associated with decreased TG levels uncovered associations with known common variation in the *BUD13-APOC3* locus, which is also associated with increased TG levels. The effects of these variants were masked by the opposite (and larger) effects of the rare variant in the Hutterites (p_gwas_=0.0002; p_conditional_=9.50×10^−9^; **Table 3**), consistent a classic epistatic interaction (**Figure S5**). The direct effects of this haplotype on the expression of the chromosome 11 apolipoprotein genes could not be assessed because their expression is restricted to the liver and had nearly undetectable levels in the LCLs.

Similarly, conditional analyses of the chromosomes 11 and 16 associations with HDL-C confirmed that the rare variant associations at these loci have independent effects from known common variants associated with HDL-C (**Table 3**). Gene expression studies of chromosome 16 variants revealed that the candidate intergenic variant located between *GOT2* and *APOOP5* associated with lower HDL-C levels is associated with higher *GOT2* expression in LCLs (p=0.004; **Figure 5C**), but showed no effect on the known lipid metabolism gene *CETP*. Overall, our results provide evidence that rare variants associated with lipid traits are likely to mediate their effects by modulating changes in gene expression, and in general have larger effects on lipid traits compared to common variants.

## Discussion

We performed GWAS with ~660k rare in European variants that occur at higher frequencies in the Hutterites, and identified four novel associations with plasma lipid traits as well as replicating the effects of the known *APOC3* splicing variant^26,27^. While the increased frequencies of these alleles in the Hutterites provided sufficient power to identify these loci in GWAS, the long stretches of LD resulted in associations with many rare variants segregating on the same haplotype and posed challenges for pinpointing the causal variant. To increase resolution and prioritize candidate variants based on their likelihood to influence each trait, we applied a statistical fine mapping approach (fgwas^28^) by jointly incorporating functional data with our GWAS results. This allowed us to narrow the subset of likely causal variants at each locus.

Three of the five rare variants identified by the GWAS in our study reside within known lipid loci identified by GWAS. However, even at those known loci, the associated rare variants in our study were independent of the known associated common variation in these regions. For example, a variant in the first intron of the *LDLR* gene (rs17242388) was associated with increased LDL-C levels in our study. Other variants in the first intron of *LDLR* have been previously implicated in regulating LDL-C levels in two studies^6,42^, but in both studies the associations had opposite effects on LDL-C levels compared to the rare variant in our study. The known variants in this intron include a predominantly European variant that is associated with lower non-HDL-C levels (total cholesterol – HDL-C) in the Icelandic population (rs17248748; MAF=3.4%; MAF = 0% in the Hutterites) and a common variant linked to multiple blood lipid traits^6,42^ (rs6511720; MAF= 15.3%; 1000 genomes EUR MAF=11.0%; p=0.009 in the Hutterites). The *LDLR* variant in the Hutterites occurs at a frequency five times higher than that reported in 1000 genomes (Europeans), and is located within a predicted enhancer region in a number of digestive tract tissues, including liver, small intestine and stomach mucosa. Conditional analyses revealed that the rare rs17242388-A allele and the common rs513663-G occur on different haplotypes and have independent and additive effects on both lowering expression of the *LDLR* gene and raising plasma LDL-C levels.

A second association with a novel rare variant also resides within a known lipid trait locus on chromosome 16q13. This is an intergenic variant (rs189679427) located between the *GOT2* gene and pseudogene *APOOP5* that is associated with lower levels of HDL-C in the Hutterites. While *GOT2* and *APOOP5* have not been directly linked to HDL-C levels, their characterized function thus far makes them interesting candidates. *GOT2* is a membrane associated fatty acid transporter that is highly expressed in the liver and several apolipoprotein genes have been implicated in lipid metabolism^22,36,37^. Moreover, rs189679427 is located 1.9Mb downstream of a well-established GWAS variant upstream of *CETP* (rs3764261; LD r^2^=0.04), a lipid metabolism gene encoding a plasma protein that supports the transport of cholesteryl esters from HDL-C to apoB-containing particles in exchange for triglycerides^43^. The rs189679427 allele was associated with lower HDL-C and with associated higher expression of *GOT2*, but not with *CETP* even though it was expressed in Hutterite LCLs. This suggests that *GOT2* may be directly involved in regulation of HDLC levels, although gene expression studies in more relevant tissues are required to confirm this observation.

The suggestive association between variants at 3q11.2 and higher LDL-C provide support for a shared genetic architecture between Mendelian and complex traits. The associated haplotype at this locus is centered around *ARL6*, a causative gene for Bardet-Bield syndrome^44^, a highly penetrant oligogenic disorder that results in a number of clinical phenotypes, including childhood obesity and hyperlipidemia in the majority of cases^45^. Many genes identified by GWAS of lipid traits also harbor loss of function mutations that underlie Mendelian disorders of lipid metabolism^37,46,47^. Mutations within these genes (coding and non-coding) provide complimentary viewpoints to the disease mechanisms influencing these traits, but further work in relevant tissues is necessary to understand the molecular basis for these associations. Overall, our results are consistent with previous findings that complex diseases are enriched for loci implicated in Mendelian traits^48^.

In summary, our findings further demonstrate the advantages of population isolates in the search for rare variants associated with complex traits. Importantly, all of the associated variation revealed in our study is within non-coding sequences and would have been missed had we focused just on exonic variation, and many were associated with gene expression, highlighting the importance of studying the effects rare non-coding variation on gene expression as well as their effects on common disease traits. An inherent limitation of this study, and most studies of rare variants, is the challenge of replicating findings due to the very low frequency of the alleles under investigation in most populations. For example, the rare and low-frequency variants surveyed in the Global Blood Lipids Consortium meta-analysis^6^ are primarily loss-of-function coding variation and had no overlap with the variants in our study, 98.9% of which were non-coding. Nonetheless, the discoveries revealed in this study, even those that may be private to Hutterites, uncovered potentially novel disease genes, and highlight new clinically relevant pathways that could point toward novel therapeutic targets for hyperlipidemias and lowering the risk for cardiovascular disease.

## Materials and Methods

### Study sample

Standard fasting plasma lipid measurements were collected as part of a larger study of complex traits in the Hutterites (see Cusanovich *et al*. (2016)^38^ for details). Briefly, blood samples were collected after an overnight fast from 828 Hutterites (ages 14 to 85 years; **Table 1**) during field trips to Hutterite colonies in 1996–1997, and 2006-2009. Plasma levels for LDL-C, HDL-C, TG and total cholesterol were measured. Subjects receiving anti-hypercholesterimia medication, hormone replacement therapy, birth control, or diagnosed with sitosterolemia^49^ were excluded from the study. For subsequent analyses, we applied a cubed root transformation to absolute LDL-C and HDL-C levels, a natural log transformation to TG and total cholesterol.

### Genotyping

We used PRIMAL^23^, an in-house pedigree-based imputation algorithm, to phase and impute 7,605,123 variants discovered in 98 whole genome sequences to 1,317 Hutterites who were previously genotyped on Affymetrix arrays^50–52^. The genotype accuracy of PRIMAL in the Hutterites was >99%, and the average genotype call rate was 87.3% due to the variation in IBD sharing across the genome of individuals with the 98 sequenced Hutterites. Within individuals, genotype accuracy was uncorrelated with call rates. See Livne *et al*. (2015)^23^ for additional details.

### Single variant and conditional analyses

We focused our studies on 660,238 variants that were rare (MAF<1%) or absent in European populations in the ExAC^53^, the ESP^54^ or 1000 genomes^55^ databases, and had genotypes called in at least 400 individuals and occurred at frequencies > 1% in the Hutterites (**Figure 1**). We refer to these variants as REVs throughout the text. To test the effect of REVs on each of the plasma lipid traits, we used a linear mixed model as implemented in GEMMA^25^ adjusting for age and sex as adding kinship as a random effect to correct for the relatedness between the individuals in our sample. Causal variants were then prioritized based on functional annotations as implemented in fgwas^28^. Follow-up conditional analyses were carried out in GEMMA for 6,781,373 variants in the Hutterites called in at least 400 individuals and with MAF> 1%. The datasets generated during and/or analyzed during the current study are available in the dbGaP repository, phs000185.

### Candidate eQTL analyses in LCLs

Candidate eQTL analyses in LCLs were performed in GEMMA and included gene expression for 441 Hutterite individuals (317 of which are in our lipid studies) that was collected as part of a separate study^38^. The LCL RNA-seq data was processed as follows. Reads were trimmed for adaptors using Cutadapt (with reads <5 bp discarded) and remapped to hg19 using STAR indexed with gencode version 19 gene annotations^56,57^. To remove mapping bias, reads were processed using the WASP mapping pipeline^58^. Gene counts were collected using HTSeq-count^59^. VerifyBamID was used to identify potential sample swaps^60^. Genes mapping to the X and Y chromosome and genes with a Counts Per Million (CPM) value of 1 (expressed with less than 20 counts in the sample with lowest sequencing depth) were removed. Limma was used to normalize and convert counts to log transformed CPM values^61^. Technical covariates that showed a significant association with any of the top principal components were regressed out (RNA integrity number and RNA concentration).

### Variant annotation

We obtained variant annotations from dbSNP, ENSEMBL, LOFTEE, conservation and functional scores (e.g. CADD, GERP, PolyPhen, SIFT), and allele frequencies from European populations (ExAC^53^, ESP^54^, 1000 genomes^55^) using Variant Effect Predictor (VEP)^62^. We downloaded promoter and enhancer annotations created by the Epigenomics Roadmap Project (−log_10_(p) ≥10; http://www.broadinstitute.org/~meuleman/reg2map/HoneyBadger2_release/) for 127 cell types or tissues. We directly annotated variants using the ClinVar database downloaded on 08/07/2016 and selected variants labelled as pathogenic or likely pathogenic. Lastly, we used 53 functional categories and 9 cell-type specific histone marks regions obtained from Finucane *et al*. (2015)^30^ (https://data.broadinstitute.org/alkesgroup/LDSCORE/). Briefly, the annotations include annotations for RefSeq, digital genomic footprint and transcription factor binding sites from ENCODE^63^, combined chromHMM and Segway annotations for six cell lines^64^, processed DHS data from ENCODE and Roadmap Epigenomics data and cell type specific H3K4me1, H3K4me3 and H3K9ac data from Roadmap Epigenomics^29^, H3K27ac from Roadmap Epigenomics and from Hnisz *et al*. (2013)^65^, super-enhancers from Hnisz *et al*. (2013)^65^, processed conserved regions in mammals from Lindblad-Toh *et al*. (2011)^66,67^ and FANTOM5 enhancers^68^. For each functional Finucane *et al*. (2015)^30^ annotation, a 500-bp window was added as an additional category. For each DHS, H3K4me1, H3K4me3, and H3K9ac sites, a 100-bp window around the ChIP-seq peak was added as an additional category.

### Fgwas

Using the fgwas software^28^ and the genomic annotations described above, we applied a single annotation model to our GWAS results to investigate enrichment of each functional categories. Similar to the procedure performed by Pickrell (2014)^28^. First, we divided the genome into ~14,000 blocks of approximately 120 Kb each (~50 variants/block) and applied forward selection to build step-wise models including the combined effects from multiple annotations. Second, we followed by a cross validation step to avoid over fitting while maximizing the likelihood of each model. We present the final best fitting models in **Figure 2**.

## Description of Supplemental Data

Supplemental Data includes six figures and five tables.

## Acknowledgements

We thank the members of the Hutterite community for their continuous participation in our studies, and the many collaborators that participated in field trips over the past 20 years. This work was supported by the NHLBI (HL085197). C.I. and S.V.M were supported by the National Institute of Health Grant T32 (GM007197) and Ruth L. Kirschstein National Research Service Awards (HL123289 and HL134315).

